# Greater host influence and promiscuity: How an invasive seaweed host has advantages over co-occurring natives

**DOI:** 10.1101/2024.12.02.626238

**Authors:** Marjan Ghotbi, Guido Bonthond, Mitra Ghotbi, Sven Künzel, David M Needham, Florian Weinberger

## Abstract

The surface microbiome of seaweed hosts is a multi-domain biofilm regulated by host-microbe and microbe-microbe interactions. The extent to which hosts influence these interactions, and potentially affect their resilience and invasion success, remains unclear. We experimentally tested whether hosts with invasion history exert more influence over their biofilms than native hosts. Biofilm formation on proxy surfaces adjacent to one invasive (*Gracilaria vermiculophylla*) and two native (*Fucus serratus, Fucus vesiculosus*) co-occurring hosts was monitored and compared to mature epiphytic biofilms of the same hosts. Only *Gracilaria’s* proxy biofilms were significantly different in community composition compared to control surfaces. *Gracilaria*’s proxy biofilms also showed the highest similarity to their adjacent algae sharing certain bacterial taxa that were absent in control treatments, indicating that colonization of the proxy surface was influenced by the host. *Gracilaria* and its proxy biofilm showed highest similarity in microbial network variables, suggesting a higher ability of the invader to influence connectivity and microbial associations within its biofilm. Meanwhile *Gracilaria*’s mature biofilm also showed higher variability in its prokaryotic composition over experiments, which was also reflected in a less robust microbial network in both *Gracilaria* and its proxy biofilms. This suggests that in addition to stronger influence in the invasive host, it was also more promiscuous towards potential symbionts from the environment. Ultimately, through examining microbial interactions, in line with previous research we found that host influence and promiscuity may play an important role in seaweed hosts to acclimate to different environmental condition and successfully thrive in new ecosystems.

## Introduction

Host-microbe and microbe-microbe interactions play a crucial role in establishing a robust microbial community of a holobiont. These interactions are affected by metabolite exchange, signalling, and physiochemical changes [1, 2]. Structural features of the seaweed microbiome as a multi-domain biofilm give it a distinctive influence on shaping these interactions. Presence of microenvironments with different osmolarity, nutrient availability, gas concentrations, and cell density of heterogeneous microbial communities stimulates the formation of three-dimensional structures within biofilm [3], which supports intercellular communication, nutrient acquisition, and protection of the microbial community [4]. While primary metabolites are recognized as inducers of microbial colonization [5], different hosts harbour distinct microbial communities [6, 7], which can occur due to host-specific signals. Some seaweeds reportedly have the ability to recruit protective bacteria and deter pathogens through secretion of surface metabolites [1]. Investigating these recruitment processes will further our understanding of how interactions between seaweeds and environmental microbiome, and also between recruited microbes determine the final biofilm composition and connectivity, and how these interactions influence resilience, dispersal and invasion success of seaweeds.

Members of the brown algal genus *Fucus* (Phaeophyta) are important habitat forming species in many shallow water regions of the northern hemisphere and especially in the Baltic Sea [8]. *Fucus vesiculosus* and *Fucus serratus* are two co-occurring species [9] in the littoral zone [8]. During the last five decades, populations of these two species in the Baltic Sea have been negatively affected by several biotic and abiotic stressors, in particular increased eutrophication, sedimentation, grazing pressure [8, 10–13], warming and seasonal variability [14]. In contrast, *Gracilaria vermiculophylla* is an invasive habitat-forming red alga (Rhodophyta) that also co-occurs with the two *Fucus* species in the Baltic Sea [15] and has successfully invaded many *Fucus* habitats despite the given threats. *G. vermiculophylla* is native to the Northwest Pacific, and has a wide invasive distribution in the Eastern Pacific [15], Eastern Atlantic including the Baltic Sea [16] and Western Atlantic [17]. There is evidence that host-microbe interactions have played a crucial role in the invasion process of *G. vermiculophylla* [1, 18]. Invasive *G. vermiculophylla* populations, compared to natives, have high host promiscuity (flexibility toward potential symbionts from the environment) [19] and exert more influence over their epibiota [20]. This enhanced influence was associated with better host performance under thermal stress, indicating it may importantly contribute to the capacity of a seaweed host to acclimate to new environments [20]. Also, co-introduction of core-microbes that provide essential functions to the host across its distribution range has been suggested to facilitate the invasion process (reviewed by [18]).

Complex host-microbe and microbe-microbe interactions within seaweed biofilms, their role in ecological success of the host, [21], and the observed shifts in the populations of the three macroalgal species, prompted us to study and compare how these hosts recruit microbes from the environment and influence microbial composition, interactions and connectivity in their biofilm. To this end, we used sterile, inert, porous artificial surfaces (polycarbonate filters of 0.22 µm pore size) in close proximity to the seaweed species without direct contact to provide substrate for biofilm formation, hereafter referred to as Proxy Biofilm (PB). This method helped to isolate the influence of exudates while eliminating other sources of variation in biofilm colonization such as host morphology [22], physical properties [23], priority effects (impact of an already established community)[24], as well as environmental factors [25, 26] which were consistent across species as they were all deployed in the same habitat. The developing proxy biofilms were then used to evaluate the hosts influence on colonization through exudates production, and were compared with mature biofilms on seaweed specimens over two experiments of different durations.

Based on the hypothesis that *G. vermiculophylla* exerts stronger host influence compared to the native *Fucus* species, we tested two sub-hypotheses; i) PBs of the invasive seaweed host are more distinct from control filters, and ii) these PBs are more similar to the host mature biofilm in terms of diversity, composition, and microbial connectivity compared to BPs of the two native hosts. We also hypothesized that *G. vermiculophylla* has higher host promiscuity, allowing more flexibility toward potential symbionts and more resilience of the biofilm to changes in the environment. Here, we also tested two sub-hypotheses; iii) the mature biofilm of invasive hosts shows short-term temporal variation in diversity and/or composition and iv) microbial connections on the invasive host and its PBs are less stable with higher propensity for reconnection compared to natives.

## Materials and Methods

### 2.1. Algae collection and experimental setup

The three seaweed species *F. serratus*, *F. vesiculosus,* and *G. vermiculophylla* were collected on 20 October 2020 from two sampling sites, Falckensteiner Strand (54°23’37.3″ N, 10°11’16.9″ E) and Bülk (54°27’15.0″ N, 10°11’50.6″ E). Six intact specimens of each species with comparable size were taken from each site. After an acclimation period of two weeks, seaweed species were cleaned from foulers, and approximately similar biomass of the species (Tab.S1-metadata) was transferred into the experimental setup. Our setup, EXUTAX (Fig.1), attached to the Kiel Outdoor Benthocosms (KOB) [27], was designed to capture the impact of host exudates (EXU) on microbial chemotaxis (TAX) and biofilm formation. We used 47 mm, 0.22 µm Nuclepore Polycarbonate Black Membrane Filters (GVS Life Sciences, Italy) as substrates for PB formation. Environmental data were collected autonomously at two-minute intervals for the appropriate depth of the setup from continuous measurements at the Kiel Fjord GEOMAR Pier (available on PANGAEA, average values reported in Tab.S1).

**Fig. 1.**
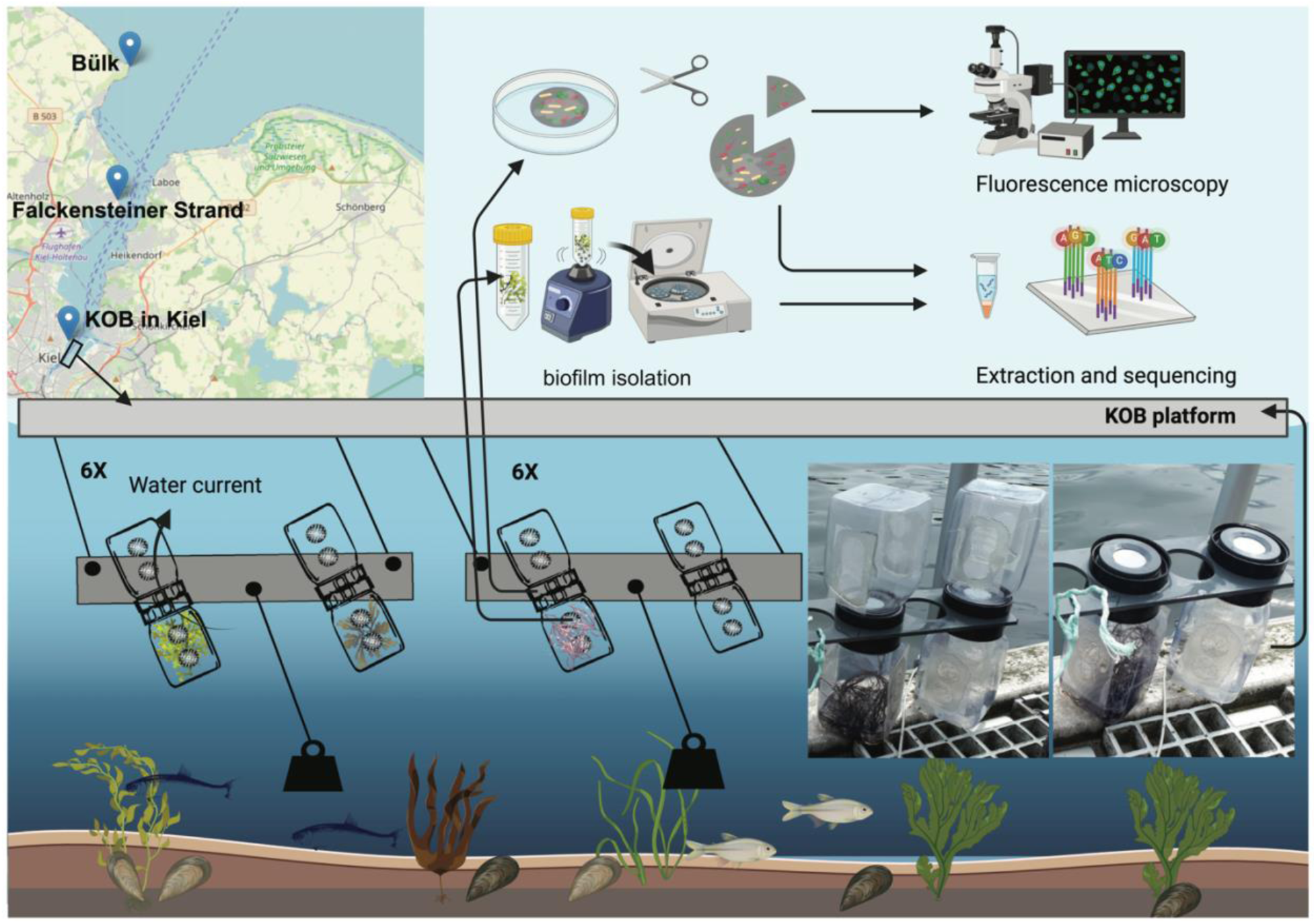
EXUTAX composed of 12 panels (plus one control panel for observation), each carrying two double-attached 1 l kautex bottles with a circular washer, as a place holder for polycarbonate filters, connecting them. Kautex bottles had four mesh-covered holes (diameter: 20 mm) on two opposing sides, allowing a current of seawater pass through constantly. The washer had a channel (diameter: 5 mm) in the middle connecting the flow between attached bottles. Seaweed specimens were deposited in individual kautex bottles, connected to an empty bottle which was harbouring the polycarbonate filter. Seaweed species and control empty bottles were arranged randomly in six replicates among panels. The whole EXUTAX was submerged in the Kiel Fjord, hanging from KOB platform, at the depth of approximately 50 cm. The set up was maintained perpendicularly by adjustment of weights to avoid sedimentation on filters. Arrangement of mesh-covered holes on two opposing sides of bottles along with the connecting channel between them allowed a current of seawater in contact with algal exudates pass through filters constantly.

### 2.2. Proxy biofilm formation, sample collection and processing

Two EXUTAX experiments were conducted in November and December 2020. In the first experiment, samples of the seaweed biofilms, PBs and control were collected on day seven (experiment spanned 3rd to 10th November), and in the second experiment on day 14 (18th November to 1st December). For the isolation of algal biofilm, approximately one g of each algal sample was taken and transferred to a tube containing sterile glass beads in 15 ml of artificial seawater. Bead beating was used to dislodge the biofilm from the seaweeds [18]. At both timepoints, 200 mL of ambient seawater was filtered on to a 0.22 µm polycarbonate filter. For studying the collected PBs, we used a combination of microscopy and molecular techniques. 1/8 of each filter was cut for enumeration via epifluorescence microscopy (supplementary information), and the rest was immediately submerged in 2% CTAB isolation buffer for DNA extraction.

### 2.3. DNA extraction and 16S rRNA gene sequencing

DNA from all biofilms was extracted following a DNA isolation Protocol for Plants [28]. The 16S-V4 region was amplified with the primers 515F (S-*-Univ-0515-a-S-19) and 806R (S-D-Arch-0786-a-A-20), as in [18, 29], and sequenced via 2 × 300 bp reads on Illumina MiSeq. Adapters were removed from raw sequence reads with cutadapt [30]. Forward and reverse reads were truncated at 220 and 200, respectively, and processed via default settings with DADA2 [31] in QIIME 2 2022.11 [32]. Amplicon sequence variants (ASVs) were classified via q2-feature-classifier [33] against the SILVA 138.1 database [34]. Subsequently, for the prokaryotic dataset, chloroplast, mitochondria, and samples less than 2000 reads were removed and gene copy numbers corrected via q2-gcn-norm (2021.04) based on rrnDB database (v. 5.7) [35, 36]. For the microalgae dataset, chloroplasts first identified by SILVA, were additionally classified with the PhytoRef database [37, 38], and samples with less than 100 reads were removed. Unassigned ASVs were reclassified when possible, using a phylogenetic approach that helps with classification of mitochondria and chloroplast (supplementary information) (Tab.S2-A; Tab.S2-B).

### 2.4. Identification of core microbiome and host influence

To evaluate host influence in attracting and maintaining specific groups of persistently associated taxa, we characterized a compositional core [39–41] at phylogenetic and short term temporal scale for both algae and PBs. ASVs with > 3 sequence reads, detected in at least 90 percent of each sample group, were considered members of the corresponding compositional core. A very low abundance threshold (0.001%) was specified to capture microbes even with rare representation in the community, since they might be physiologically more active compared with more abundant ones [39, 42]. In addition, to capture host specificity, we detected ASVs present in 100% of each seaweed species and their corresponding PBs across both experiments, but absent in control filters and seawater samples.

### 2.5. Microbial association network construction

To evaluate cross-domain associations (significant abundance correlation between and within prokaryotes and microalgae), SPIEC-EASI network analyses [43] were applied at both the whole microbial community (WMC) and core microbiome (section 2.4) levels on merged ASV datasets of seaweeds, and PBs including control samples (supplementary information). For WMC, ASVs with < 3 reads and < 0.25% minimum relative abundance were excluded (this abundance threshold was used since it was the minimum required for *Fucus* hosts networks to reach stability). Hub taxa, module (a cluster of interconnected microbes) and network hubs, were identified for each sample group based on within- and among-module connectivity [44, 45].

### 2.6. Statistical analysis

#### 2.6.1. Diversity and quantification

Linear mixed models (LMM) [46] from the R package lme4 were used to estimate the comparative and interactive effects of seaweed exudates on prokaryotic and microalgal diversity (Shannon indices) on both living (mature biofilm) and non-living (polycarbonate filter) substrates across two experiments. LMM was also applied to evaluate the impact of seaweed exudates on the enumerated values of microalgae. Seaweed exudate impact on mature biofilm (n=33) and PBs (n=35) and timepoint were considered as fixed factors. The factor panel was included as random effect to account for non-independence among observations from bottles installed on the same panel and the factor bottle to account for non-independence between filters and seaweeds from the same bottle over two experiments.

#### 2.6.2. Community composition

Variability in microbial composition was analysed using permutational analysis of variance (PERMANOVA) with Bray-Curtis distance matrices, after testing for homogeneity of dispersion (beta dispersion), through vegan package [47] with 999 permutations. ANCOM-BC [48] was used to find differentially abundant ASVs (at phylum level) in *G. vermiculophylla* mature biofilm between two experiments.

#### 2.6.3. Microbial connectivity

To identify similarity between microbial connectivity in distinct biofilms, we generated a Euclidean distance-based analysis using mean values of network variables calculated from significant connections detected in each network which visualized through PCA and heatmap.

## Results

### 3.1. Microbial community composition across seaweed biofilms and PBs

The prokaryotic community analysis, after processing, had 12,685 ASVs and 2,431,184 reads across 87 samples (36 seaweeds, 35 PBs and 14 control filters, and two ambient water). Seaweeds had the highest number of unique ASVs (2655) followed by PBs and control filters (1074) and 179 ASVs were shared between all samples (Fig.S1-A). The microalgae dataset (via chloroplasts) had 187 ASVs and 827,700 reads across 84 samples (33 seaweeds, 35 PBs and 14 controls, and two ambient water). PBs and control filters had the highest number of unique microalgal ASVs (73) vs. seaweeds (18) and 25 were shared between all (Fig.S1-B).

Across all surfaces (seaweeds, PBs, and control filters), Proteobacteria was the most abundant phylum, followed by Bacteroidota and Planctomycetota. However, in ambient water, Crenarchaeota was the second most abundant phylum after Proteobacteria (Fig.S2-A). Desulfobacterota and Campylobacterota phyla were mainly associated with seaweed samples, with higher abundance on *G. vermiculophylla*. Firmicutes and LCP-89 showed high relative abundance exclusively in the *G. vermiculophylla* biofilm, while Spirochaetota and Acidobacteriota were more abundant in *Fucus* species. The microalgae community on all surfaces was strongly dominated by Bacillariophyta, while in ambient water it consisted of Bacillariophyta and Cryptophyceae (Fig.S2-B).

Analysis of the 30 most abundant prokaryotic ASVs showed higher abundances of *Spongibacter, Neptomonas* and *Desulforhopalus* on the *G. vermiculophylla* biofilm and its PBs (Fig.2-A). *Fucus* species and their PBs showed higher abundance of ASVs from the BD1-7 clade (specifically ASV1404), however, *G. vermiculophylla* biofilm and its PBs were almost devoid of this ASV (Fig.2-A). Generally, PBs and control filters were enriched with Alphaproteobacteria. Among microalgae, ASVs belonging to Melosiraceae, Coscinodiscophyceae, Cymbellaceae, Bacillariophyceae were strongly associated with surfaces. However, Thalassiosirales and Pyrenomonadales were mainly found in ambient water.

**Fig. 2.**
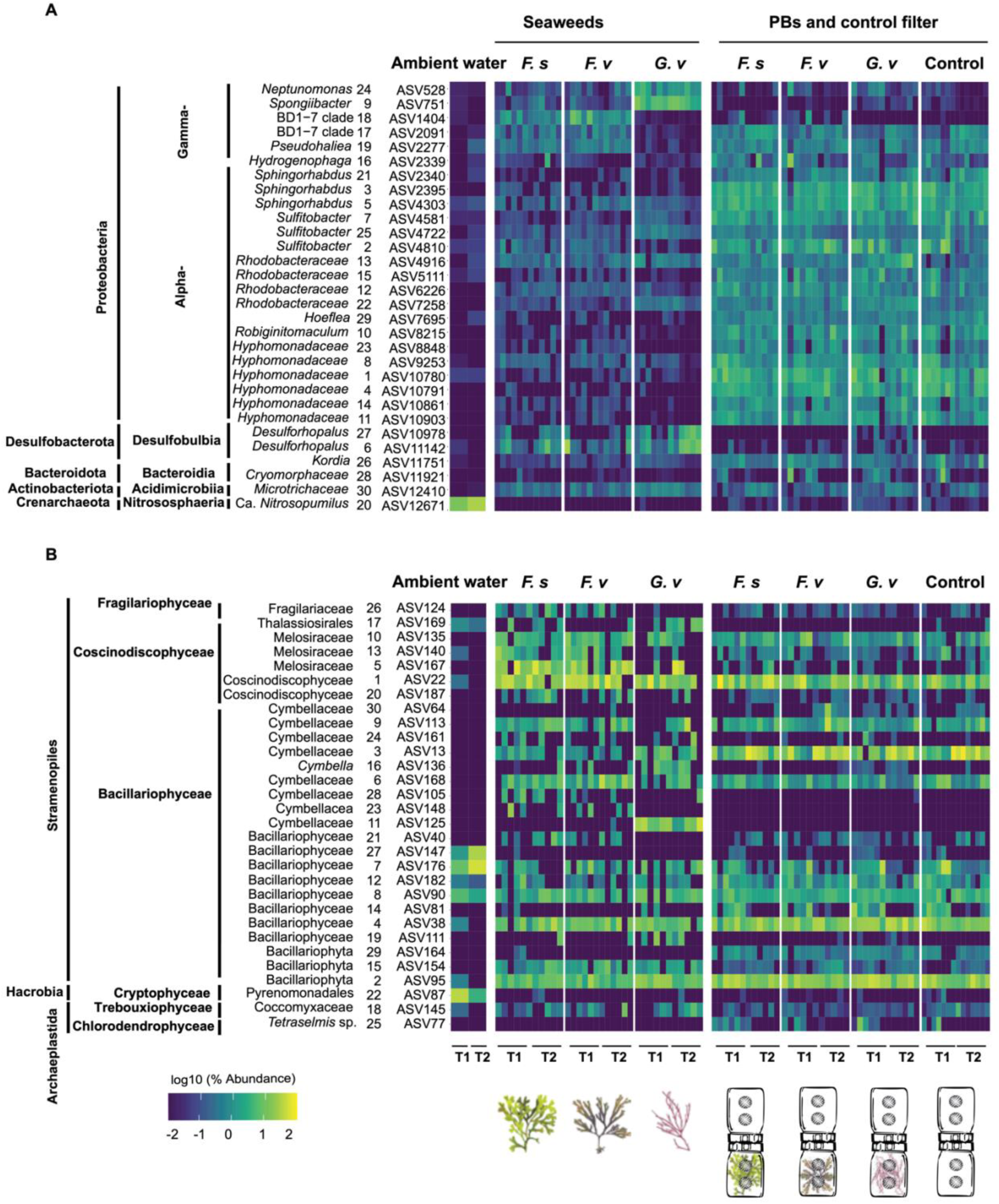
Heatmap representing the top 30 most abundant A) Prokaryotic and B) Microalgae ASVs during two experiments. Each sample group (seaweeds, PBs and control) has six replicates in each timepoint except for ambient seawater with two replicates and controls with one additional replicate). ASVs are classified at their highest detected resolution. The numbers printed on the lefthand side of ASV codes represents their order in relative abundance, with 1 being the most abundant ASV. The relative abundance of ASVs is log10 transformed. Seaweed and filter sample illustrations are shown below the heatmap for reference.

Some members of the Cymbellaceae family showed degrees of host specificity. For instance, ASV125 was consistently associated with *G. vermiculophylla* and not detected among abundant ASVs of other samples, while ASV81 and ASV154 were primarily detected on *Fucus* species and not among the abundant ASVs of PBs (Fig.2-B).

### 3.2. Microbial profiling and quantification

In both experiments, prokaryotic diversity (Shannon indices) exhibited no significant differences between control filters and *G. vermiculophylla*’s PBs, and generally diversity stayed similar across all treatments on non-living substrate (Fig.3-A; Tab.S3-B). Only diversity of *G. vermiculophylla*’s PBs was similar to their corresponding algal treatment and native hosts showed significant differences with their PBs (Fig.3-B). The highest prokaryotic diversity was observed in the mature biofilm of *Fucus* algae, and their Shannon indices were significantly higher compared to *G. vermiculophylla* and to PBs (Fig.3-B). While diversity on non-living substrates was significantly higher during the second (longer) experiment (Tab.S3-B), no significant change over time was observed on mature algae biofilms (Tab.S3-C; Fig.3-C). Likewise, microalgae diversity also stayed similar on non-living substrates across all algal treatments, and no significant differences were detected between control filters and *G. vermiculophylla*’s PBs (Fig.3-D). All algae and their PBs showed similar diversity over both experiments (Fig.3-E) and the only difference was seen among substrate types, primarily in the second experiment (Tab.S3-D). Enumeration of microalgae observed on PBs and control filters (n=47) except for between two experiments, did not show any significant difference between algal treatments and control (Fig.S3; Tab.S4).

**Fig. 3.**
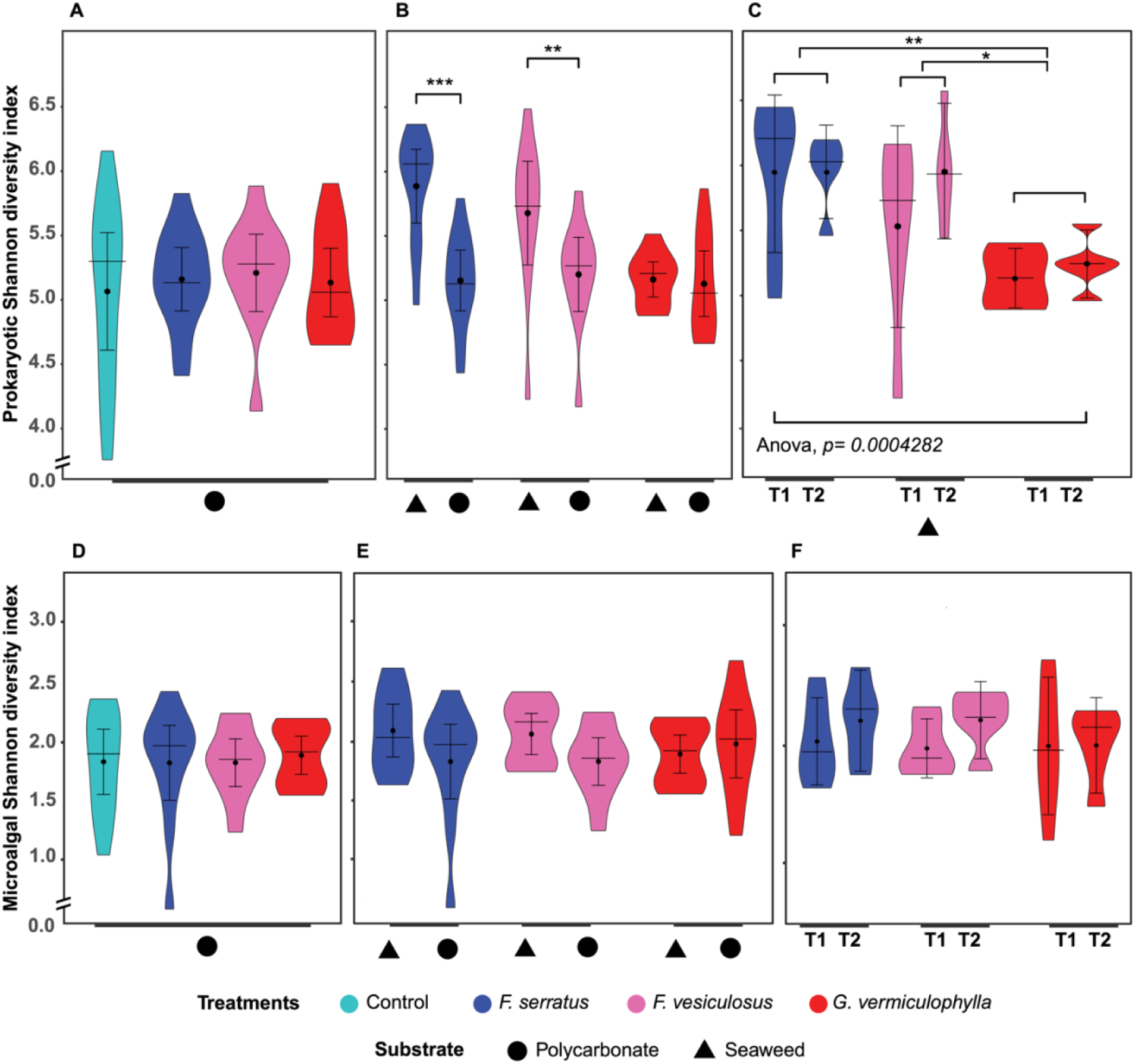
Diversity (α-diversity) of A) Prokaryotes and B) Microalgae for different samples on living (seaweeds) and non-living (polycarbonate) substrate types over two experiments. Violin plots showing median, interquartile range (with outliers) and the point and the bar showing the mean and the standard deviation. Six replicates at each timepoint were used for different treatments.

The prokaryotic community on seaweeds biofilm was fully separated by PCoA from that on PBs along the first principal component axis PCoA1, while PCoA2 mainly separated the biofilm of two algae genera (Fig.4-A). The PERMANOVA likewise detected significant impact of living and non-living substrate (seaweed vs. PBs; R^2^=0.18, *p*=0.0001; Tab.S5-A). The prokaryotic community on non-living substrate (n=49) showed significant impact of algal treatments. Pairwise ADONIS detected only a significant difference between PBs of *G. vermiculophylla* and control filters (R2=0.06, *p*=0.048; Tab.S5-B). Pairwise ADONIS on algae mature biofilms (n=36) showed a significant difference of *G. vermiculophylla* and the two *Fucus* species (Tab.S5-C). All developing biofilms on non-living substrates and only *G. vermiculophylla’*s mature biofilm (n=12) exhibited significant shifts of prokaryotic communities over time as an influence of environmental changes (Tab.S5-B,C). Observed environmental data over the two experiments showed differences in salinity, temperature, oxygen saturation levels and irradiation, with temperature in *G. vermiculophylla’*s BP and mature biofilm as well as salinity (conductivity) in all non-living substrate samples being the main source of variation (Tab.S.6-A:C). Microalgal communities also mainly differed between substrates, as indicated by the separation along PCoA1 (Fig.4-B), and detected by PERMANOVA (n=82; R^2^=0.19, *p*=0.0001; Fig.4-B; Tab.S5-D). The microalgal communities on seaweed surfaces (n=33) showed a significant difference between *G. vermiculophylla* and the two *Fucus* species (Tab.S5-F), consistent with patterns observed in the prokaryotic community. Microalgae showed a significant shift over time on both substrates suggesting an impact of environmental factors (Tab.S5). In contrast to the prokaryotic community, the presence of different seaweed treatments had no significant effect on the microalgal community on non-living substrates (Tab.S5-E).

**Fig. 4.**
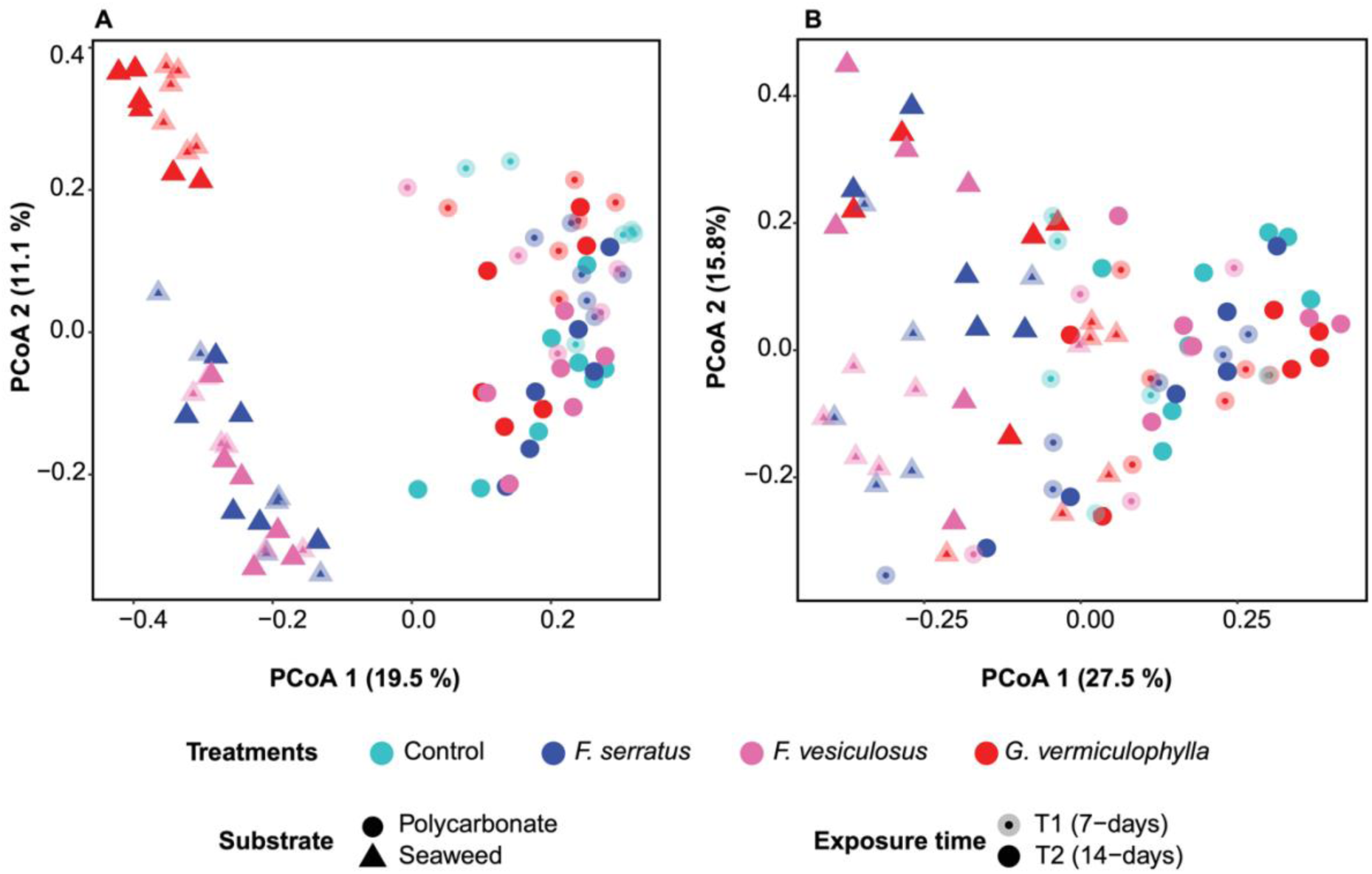
PCoA plots demonstrating A) Prokaryotes and B) Microalgae community clustering based on Bray-Curtis dissimilarity matrices. The number of replicates for sample groups for each experiment was six, except for controls with seven replicates.

### 3.3. Core microbiome

*G. vermiculophylla*’s biofilm showed the highest ASV and phylum level diversity in its prokaryotic core microbiome, with 175 ASVs vs. 148 for *F. vesiculosus*, and 94 for *F. serratus* (Fig.S4). Members of two *Fucus* species shared a higher number of their core ASVs compared to *G. vermiculophylla.* Core taxa of PBs compared to seaweeds showed higher diversity at ASV but lower at phylum level. PBs showed the same trend among seaweed species with 191, 189 and 137 ASVs correspondingly. Controls possessed the lowest diversity of ASVs (93) and phyla within their core (Fig.S4; Tab.S7-A). Although the differences were not statistically significant, they highlight *G. vermiculophylla*’s potential for maintaining a broader range of prokaryotic diversity in its core taxa. The microalgae core taxa were generally lower in numbers compared to prokaryotes (Fig.S4). Likewise, PBs showed higher diversity of ASVs compared to seaweeds (Fig.S4; Tab.S7-B). Two prokaryotic ASVs, identified as Rhodobacteraceae and *Granulosicoccus,* were present in 100% of the *G. vermiculophylla*’s biofilm and its PBs, but were rarely and in very low abundances detected in other seaweed samples and absent from other proxy biofilms. Presence of these host-specific ASVs denotes higher similarity between *G. vermiculophylla*’s biofilm and PB, suggesting stronger influence of *G. vermiculophylla*’s exudates in attraction of specific bacteria in the environment (Tab.S7-A).

### 3.4. Microbial association network

The WMC of *Fucus* species showed the most similarity (Fig.5-A) and were strongly separated from *G. vermiculophylla* and PBs, consistent with diversity results. PBs were distinct from control biofilms mainly in core microbiomes (Fig.5-A,B). The most dissimilarity between PBs and control filters was seen in the core microbiome of *G. vermiculophylla*. In terms of microbial connectivity, the most similarity between seaweeds and their PBs was seen in *G. vermiculophylla* for both WMC and core microbiome (Fig.5-A,B). Conversely, the most dissimilarity was seen in *Fucus* species and their PBs in both WMC and core microbiome (Fig.5-A,B).

**Fig. 5.**
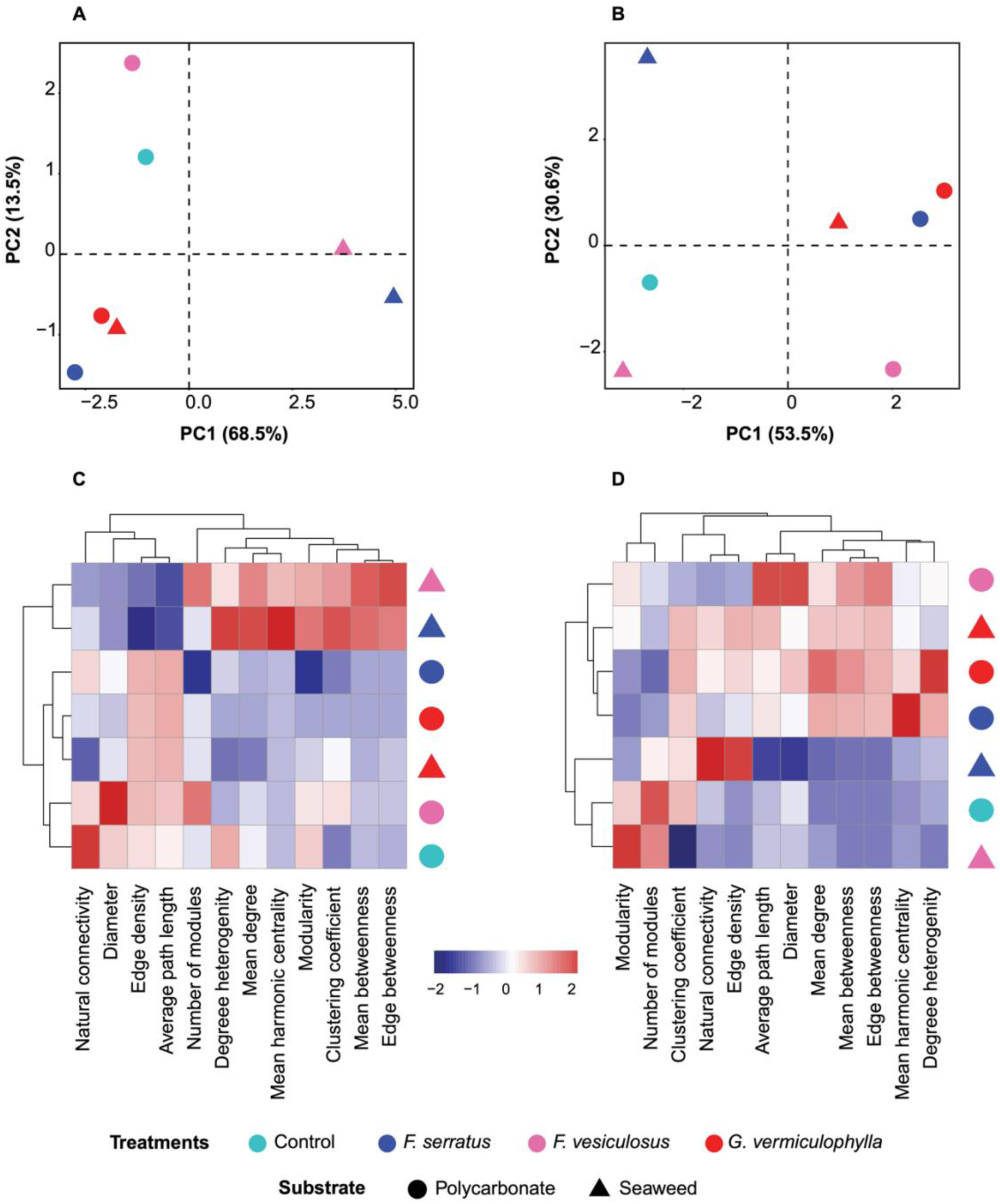
PCA plot where distances represent the dissimilarity between microbial association networks of A) whole microbial community (WMC: > 0.25% relative abundance) and B) core microbiomes (≥ 90 prevalence) of seaweed biofilms and their PBs and control. Heatmaps comparing network variables for C) WMC vs. D) core microbiomes of the corresponding samples.

Beyond the broad overview of similarities, in terms of various individual metrics, microbial networks showed opposite trends at WMC vs. core microbiome of each seaweed (Fig.5-C,D). First, among WMCs, the sparsest network was seen in *G. vermiculophylla* mature biofilm with the lowest average number of connections between nodes or ASVs (mean degree) compared to *Fucus* species (Fig.5-C; Tab.S8). However, between core microbiomes, *G. vermiculophylla* and its PBs had among the highest number of connections, along with *F. serratus* PBs (Fig.5-D). Relatedly, the robustness of the networks as a function of natural connectivity [49], was lowest for the WMC of *G. vermiculophylla* mature biofilm; however, its core microbiomes of both mature biofilm and PBs showed among the highest corresponding values after *F. serratus* mature biofilm (Fig.5-C,D). The clustering coefficient (fraction of observed vs. possible node clusters) which represents the complexity of the network due to strong interactions among microorganisms [50, 51], also showed the same trend between the *G. vermiculophylla’s* WMC to its core microbiome, with *G. vermiculophylla’*s core on both mature biofilm and PBs showing the highest network complexity (Fig.5-C,D; Fig.6). The highest modularity, denoting denser connections between the nodes within modules but sparse connections between nodes of different modules [45, 51], was observed in the WMC of *Fucus* species than *G. vermiculophylla* (Fig.5-C). While the highest modularity of WMC in algae mature biofilm was with *F. serratus*, its core microbiome showed the lowest value (Fig.5-C), which denotes presence of less stable niches in its core taxa. Among core microbiomes, frequencies of associations and specifically negative associations were significantly lower in *Fucus* species (Fig.6).

**Fig. 6.**
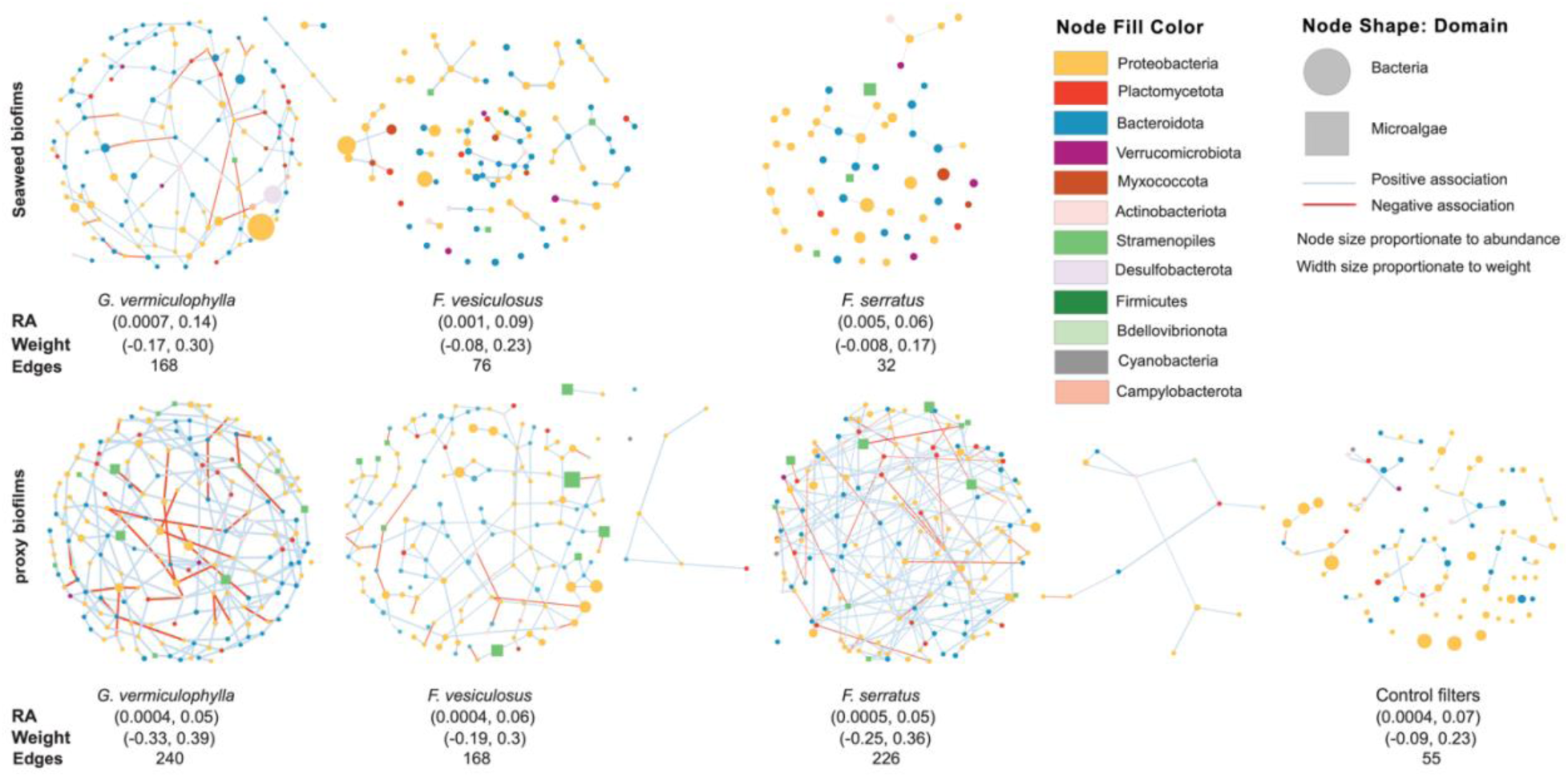
Microbial association networks of core microbiomes of algae and PBs including bacteria and microalgae. Each node represents an ASV and is shaped according to the taxonomic domain. Edge color denotes a positive (blue) or negative (red) association between two connected ASVs with the width proportional to weight (correlation coefficient of nodes abundances representing strength of associations). The corresponding maximum and minimum values for RA (Relative Abundance) and weight, and number of edges are provided underneath each network plot. Generally, the highest number of associations are seen in the networks of *G. vermiculophylla* among both seaweeds and PBs.

Regarding the taxonomic membership, the networks of WMC of seaweeds biofilms consisted of nodes from three microbial domains including prokaryotes (bacteria, archaea) and eukaryotes (microalgae). Regarding bacteria, *G. vermiculophylla* possessed the highest number of nodes from *Desulfobacterota*, with the *Desulforhopalus* accounting for the highest degree (number of connections) and betweenness (centrality). However, *Fucus* species possessed higher abundance and diversity of Proteobacteria, Bacteroidota and Planctomycetota. For archaea, *Fucus* species as well as PBs had *Nitrosopumilus* (shared with ambient water) (Fig.S5-A:B), while *G. vermiculophylla* possessed two archaeal ASVs as *Methanolobus*, a psychrophilic methanogen and SCGC_AAA286-E23 (candidate phylum Woesearchaeota) (Fig.S5-C; Tab.S9). SCGC AAA286-E23 is reported as anaerobic [52] symbiont of other prokaryotes [52–54]. Higher among-module connectivity of this archaeon, as a connector node (a node connecting within and between microbial sub communities [45]), supports its symbiotic lifestyle (Tab.S9). In all seaweed’s mature biofilm, dominant contributors of microalgae community were Stramenopiles (Bacillariophyta). *G. vermiculophylla* in addition, had a representative from Archaeplastida (Coccomyxaceae) with relatively high betweenness and degree (Fig.S5-C; Tab.S9). ASV125 (Cymbellaceae) which was only detected in high abundance on *G. vermiculophylla* (Fig.2-B) and classified as a connector node (Tab.S9) was in the same module with *Methanolobus* and sulfur cycle bacteria (Fig.S5-C), suggesting potential metabolic interactions among these microbes.

In the network of core microbes, 46 ASVs and three associations were shared between *G. vermiculophylla* and its PBs, five of which were not detected on control filters (Tab.S10). In particular these three associations were between Rhodobacteraceace with *Eudoraea*, *Sulfitobacter* with *Sulfitobacter*, and Hyphomonadaceae with Flavobacteriaceae (Fig.7). *F. vesiculosus* and *F. serratus* shared 50 and 26 ASVs with their PBs networks, but no associations (Tab.S10). However, with the exception of two ASVs in *F. vesiculosus* all were shared with control filters (Tab.S10). To examine the individual taxa that could have outsized roles in maintaining connectivity and functioning of the network, we analyzed ‘hub taxa’, which are a small number of strongly interconnected microbes [45, 55]. Hub taxa for different seaweeds and PBs were defined based on within-module and among-module connectivity of nodes within each network [44, 56]. *Sulfitobacter* (Alphaproteobacteria), and P3Ob-42 (Myxococcota) were identified as hub taxa for *G. vermiculophylla.* OM190, *Phycisphaera*, *Blastopirellula* (Planctomycetota), *Lutimonas* (Bacteroidota), and *Nannocystis* (Myxococcota) composed hub taxa of *F. vesiculosus*. *F. serratus* hosted ASVs from Rhodobacteraceae (Alphaproteobacteria), *Colwellia* (Gammaproteobacteria), Arcobacteraceae (Campylobacterota), Planctomicrobium (Planctomycetota), Cryomorphaceae, Saprospiraceae, Wenyingzhuangia (Bacteroidetes) as its hub taxa (Fig.S7; Tab.S11).

**Fig. 7.**
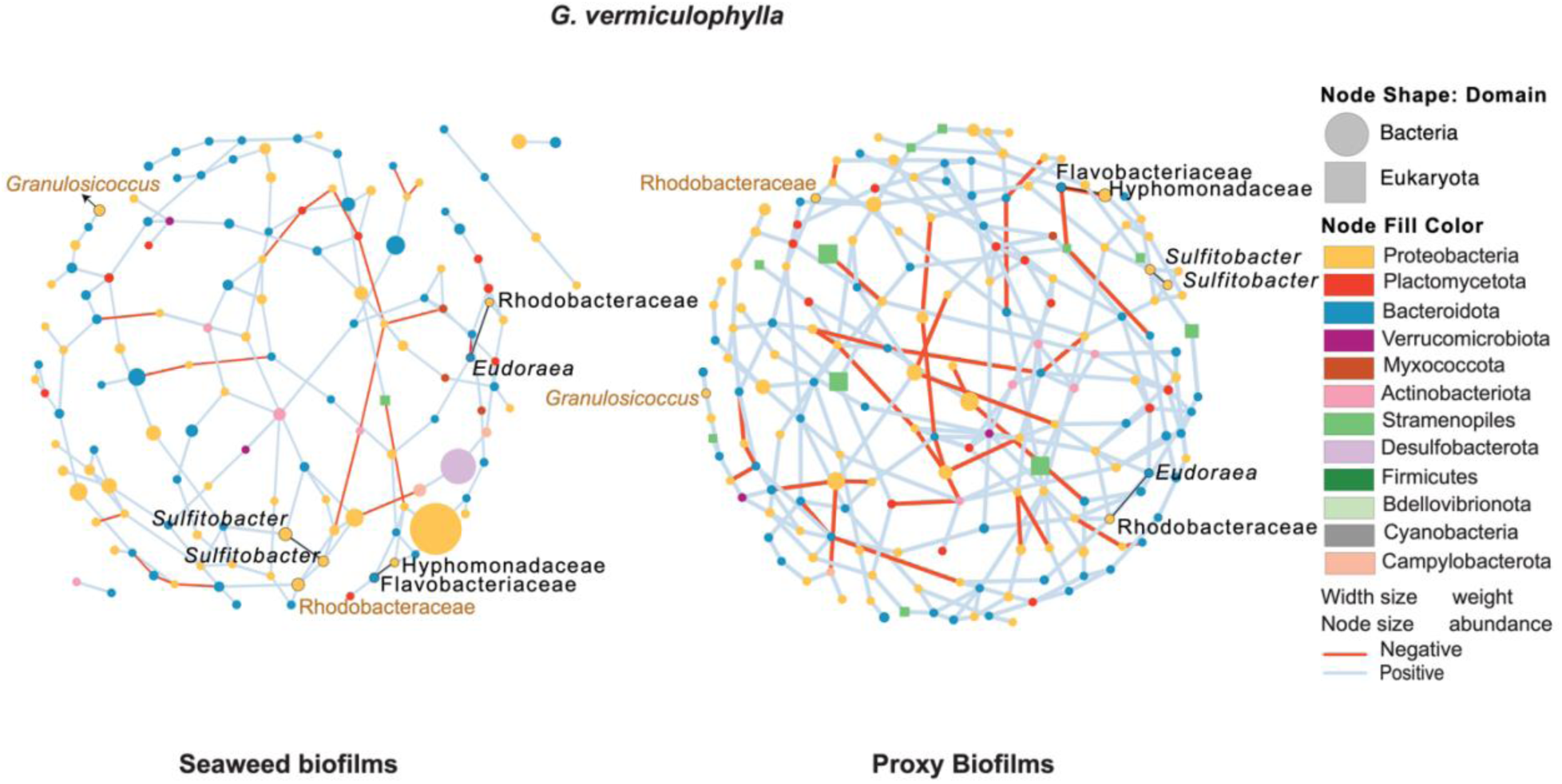
Comparison of microbial association networks in *G. vermiculophylla* seaweed and its PB with 46 nodes (ASVs) and three associations (edges in black) shared between them. Node shape represents microbial domain and node color shows their phyla. Edge color indicates a positive (blue) or negative (red) association and the edge width is proportional to weight. Two ASVs in brown color are hosts-specific taxa exclusively shared between *G. vermiculophylla* and its PBs.

## Discussion

### 4.1. Host influence on microbial composition and interactions in its biofilm

The ability of hosts to influence the biofilm composition can potentially aid their dispersal across diverse environments, contributing to invasion success [20]. Using an *in-situ* approach, we found that the invasive alga (*G. vermiculophylla*) expressed more host influence over its biofilm and more promiscuity toward potential symbionts and their interactions, compared to the native species (*F. serratus and F. vesiculosus*). *G. vermiculophylla* caused significant difference between its PBs and control filters composition (Tab.S5-B). This invasive alga also showed higher similarity between its mature biofilm and PBs regarding terms of diversity (Tab.S3-A), and microbial connectivity (Fig.5) which suggests that the host influenced the microbial community on the PBs at a structural level. Further, more microbial taxa, including host-specific taxa (Fig.7) were shared between *G. vermiculophylla* and its PBs, compared to *Fucus* species. This supports the first and second hypotheses, highlighting the invasive host’s stronger host influence on its PBs. This influence is likely driven by host-produced exudates, exchanged through the PB filters, which may attract and/or deter environmental microbes [1]. A positive association between Hyphomonadaceae and Flavobacteriaceae was seen on *G. vermiculophylla*. Both families have been recurringly reported on *G. vermiculophylla* and other macro [57–59] and microalgae [60]. Additionally, two ASVs of Rhodobacteraceae and *Granulosicoccus* showed high degrees of host specificity and were exclusively shared between *G. vermiculophylla* and its PBs. These taxa were reported to be among winter core microbiome of *G. vermiculophylla* [59]. Members of the Rhodobacteraceae family are known for association with initial surfaces colonization in marine ecosystems due to their ability to react to low levels of nutrients faster than other bacteria [61]. Hence these bacteria may be among the initial colonizers of *G. vermiculophylla* with influence on community succession. *Granulosicoccus*, a flagellated bacterium capable of chemotaxis [62], is one of the rare Gammaproteobacteria that encode a DMSP demethylase, widely found in marine Alphaproteobacteria [62, 63]. Epiphytic *Granulosicoccus* also has been reported to carry metabolic genes for nitrate and nitrite reduction [62], sulfur transformation [64], and vitamin B-12 production [64], suggesting critical roles for this genus in nutrient cycling and vitamin acquisition for both the auxotrophic host and its biofilm assembly.

### 4.2. Distinct biofilm connectivity and microbial associations on invasive and native hosts

Invasive and native seaweeds supported distinct microbial diversity and communities. Their networks of association also differed between WMC and core microbiomes. Algal PBs had similar communities and diversity but differed in microbial association networks. The WMC of *G. vermiculophylla* mature biofilm demonstrated the lowest robustness (lower mean degree and natural connectivity) and at the same time the highest density of connections (number of present connections to all possible connections) which can potentially denote flexible reassembly and thus high host promiscuity, which supports our third and fourth sub-hypothesis that *G. vermiculophyllaa* is more promiscuous toward potential symbionts. This host trait may importantly promote acclimation to changing environmental conditions. This was also supported by the networks of *G. vermiculophylla*’s PBs which revealed less stable microbe-microbe connectivity compared to *Fucus* BPs (Fig.5), suggesting the invasive host is less dependent on the WMC community. In contrast to WMC, the core microbiome of *G. vermiculophylla*, on both mature biofilm and PBs, possessed a higher number of connections (edge density), and showed stronger associations and more developed and complex connectivity compared to core taxa of *Fucus* species (Fig.6). This suggest that while *G. vermiculophylla’*s biofilm is generally flexible toward environmental microbes, it persistently maintains a diverse and interconnected core community. Unlike *G. vermiculophylla*’s WMC, its core microbiome exhibited a higher clustering coefficient in both mature biofilm and PBs (Fig.5-C,D), indicating a greater potential for niche differentiation. This theoretically facilitates coexistence of diverse core microbial taxa by reducing direct competition [65], which aligns with our findings on higher diversity in *G. vermiculophylla*’s core taxa (Tab.S7-A). Presence of functionally developed microbial subcommunities within the core microbiome of *G. vermiculophylla* may aid with nutrient acquisition and defence. Contrarily, the *F. serratus* biofilm with the highest prokaryotic diversity in its WMC revealed the most depauperate core microbiome. *F. serratus* core taxa, while displaying the highest connectivity relative to their number, showed the weakest associations (Fig.5-D; Fig.6). Generally, *Fucus* species and specifically *F. serratus,* showed weaker microbe-microbe associations within their core microbiome niches (Fig.6). This denotes less stability of functional groups within *Fucus* species core microbial taxa.

Networks of *G. vermiculophylla*’s biofilm, both in WMC and core, possessed the highest number and diversity of *Desulfobacterota,* the largest phylum harboring sulfate-reducing bacteria (SRB) [66], with *Desulforhopalus* ASVs having an intermediary role in facilitating connections between other taxa. Members of the Desulfocapsaceae family exhibit diverse phenotypic characteristics, such as a wide temperature tolerance, different motility properties, anaerobic chemolithotrophic or chemoheterotrophic metabolism, and utilization of various electron donors and acceptors for sulfate reduction, which helps them to thrive under different environmental conditions [66]. Presence of this taxonomic guild in high abundance can contribute to resilience of the host biofilm and its function under different environmental conditions. Seaweed hosts and their associated microalgal community usually provide a substantial quantity of organosulfur compounds that can be degraded by bacteria [67]. Numerous bacteria produce extracellular enzymes that require suboxic or anoxic environments to support microaerophilic or anaerobic metabolism. Anaerobic microniches formed in biofilms could facilitate this [68], and enhance nutrient cycling that benefits their hosts. An increase in the relative abundance of certain anaerobic prokaryotes including symbiotic Woesearchaeota [52–54], along with aerotolerant anaerobic Cloacimonadota [69] in the *G. vermiculophylla* biofilm in the second experiment (Fig.S6) implies the presence of microniches influenced by host rather than direct impact of environment. Sulfate availability in anoxic environments can facilitate DMS degradation by methanogens and SRB [70]. The positive association between sulfur cycling bacteria and methanogenic archaea (Fig.S5-C), suggests presence of such syntrophic relationship between these groups within the *G. vermiculophylla* biofilm similar to what has been observed in anoxic sediments [70].

Hub taxa, have been reported to be able to reflect controlling impact of the host genotype on microbiome assembly [55]. In short, the environment directly affects “hub” microbes, and this effect transmits to the microbial community via microbe–microbe interactions [55]. P3Ob-42 (Myxococcota phylum) was identified as most central hub taxon of *G. vermiculophylla* with the highest connectivity within and between subcommunities (Tab.S11; Fig.S7). P3Ob-42 is a potential sulfate reducer and methane oxidizer and contributes to nitrogen and phosphate cycling [71]. It is reported to be associated with good health status of marine hosts, specifically Carrageenophyte red algae and corals [72, 73]. *Sulfitobacter*, the second hub taxon of *G. vermiculophylla* (Fig.S7), is known for its potential DMSP degradation, growth promoting impact and contribution to the health of algae and corals in cold marine environments [72, 74, 75]. Hub taxa in *F. serratus* included Planctomycetota, Bacteroidota, and Campylobacterota. *F. vesiculosus* also included Bacteroidota but was dominated by hub taxa from Planctomycetota (Fig.S7). *Fucus* spp. secrete fucoidan, a sulfated polysaccharide, containing high percentages of L-fucose and sulfate ester groups [76], that serve as a substrate for the abundant sulfatases produced by Planctomycetota and favour their colonization [77]. In addition, The peptidoglycan-free cell wall of most Planctomycetota enables resistance to antimicrobial activities from host and other bacteria in the biofilm [77].

### 4.3. Greater acclimation potential of invasive host’s biofilm to environmental changes

Prokaryotic community composition in *G. vermiculophylla’s* mature biofilm showed a significant shift over time (Tab.S5-C). This shift may be driven by environmental changes, including water temperature, salinity, oxygen levels and irradiation impacting host [78] and hence its biofilm during the second experiment (Tab.S1; Tab.S6-A). This observation is an additional support to the third hypothesis that the invasive holobiont is more promiscuous and can undergo greater shifts in its biofilm composition over time [19]. The host promiscuity was also supported by less robust microbial networks in both *G. vermiculophylla* and its proxy biofilm. Host promiscuity may enable a host to associate a taxonomically or compositionally different microbiome, which maintains functions essential to the host [79]. This may be an important trait for seaweeds, facilitating acclimation and potentially supporting biological invasions [19]. Whereas previous work on *G. vermiculophylla* showed that host promiscuity varies between native and invasive populations of the same species [19], this study provides evidence that host promiscuity also varies between invasive and native species co-existing in the same environment that can impact their competitions and ecological success.

## 5. Conclusion

While previous work has found that invasive *G. vermiculophylla* populations have greater host influence [20] and promiscuity than its native populations [19], this study is the first to compare invasive versus native hosts of different species coexisting in the same environment using an *in-situ* experiment isolating the impact of exudates from the host substrate itself. It is also the first study to look specifically at microbial interactions of the given hosts, providing a new layer of evidence on how host influence and promiscuity at the level of microbe-microbe interactions may drive seaweeds invasions. In addition, we found stronger host-microbe and microbe-microbe associations within a set of conserved core microbes associated with *G. vermiculophylla*, despite its higher host promiscuity. This suggests that some taxa fulfil key functions and are not easily replaced, and may have accompanied their host through the invasion process. Although some of these taxa were recruited in the new environment, further research is needed to unravel the native or invasive origin of these specific symbionts. Ultimately, our results suggest that host influence plays an important role in seaweed holobionts. While the exact identity of exudates remains unknown, this study demonstrates that seaweeds manipulate the composition and connectivity of microbial communities in their proximity. Future study is needed to characterize the metabolites through which the host achieves this and develop a mechanistic understanding of how seaweed biofilm is shaped by their host.

## DATA AVAILABILITY

The raw de-multiplexed V4-16S rRNA gene reads and corresponding metadata were deposited in the SRA database under the BioProject accession number (PRJNA1180617). Environmental data for temperature and salinity during the given period is available in PANGAEA (https://doi.org/10.1594/PANGAEA.963281) [80], and the average values of other environmental data (irradiation, oxygen, salinity) are available in the metadata file (Tab.S1). Scripts for analysis and figures are available via https://github.com/Marjan-Ghotbi/Seaweeds-Microbial-dynamics.

## AUTHOR CONTRIBUTIONS

MaG and FW conceptualized the study. Field collection, experiments, and laboratory work were conducted by MaG, with supervision from FW. Data processing and analysis were carried out by MaG, with supervision from DMN, and FW. MiG and GB provided valuable input regarding statistical analysis. All authors contributed to the writing and revision of the manuscript.

## ACKNOWLEDGEMENTS

This study was funded in part by GEOMAR institutional funding received by Martin Wahl and FW, who supported the setup construction, supplies, and materials. Further support was provided by a Young Investigator grant awarded to DMN. We are especially grateful to Martin Wahl for his instrumental role in the experimental design and conceptualization of the study. Our thanks also go to Nadja Stärck and Björn Buchholz for their invaluable assistance during sample collection and setup construction.

