## Supplementary information for "Greater host influence and promiscuity: How an invasive seaweed host has advantages over co-occurring natives"

### Materials and methods

#### Sample collection and processing

To avoid pseudoreplication of seaweeds, we collected individuals that were attached to hard substratum to ensure not sampling fragments separated from the same individual or clonally reproduced drifting individuals and deposited them in the EXUTAX set-up. For biofilm sample collection at the end of each experiment, all panels were taken out of water at approximately the same time to terminate the incubation period equally for all treatments. For sampling of seaweeds' biofilm, approximately one g of the thallus of each algae specimen was cut and stored in 50 ml sterile tubes. All samples were immediately stored in a cooler until arrival at the laboratory, and transferred to a 50 ml tube containing 10 sterile glass beads of 4 mm diameter in 15 ml of artificial seawater. To collect the epiphytic microbiome, tubes containing glass beads were vortexed for 5 min, then the suspension was spined down for 15 minutes [1]. The precipitated microbiome was preserved in ethanol and stored at -20. Potentially some firmly attached epiphytic microbes may remain associated to the algae surface despite this vigorous effort to dislodge them. The resulting data are included in the metadata (Tab.S1).

Between each time of filtration for ambient water samples, pieces of vacuum pump in contact with sample were washed with sodium hypochloride to remove any DNA cross contamination, then rinsed 3 times with sterile distilled water to eliminate sodium hypochlorite (NaClO) and avoid target DNA destruction. Filters were stored in - 20 °C for molecular identification.

Temperature, salinity and oxygen data were logged in with two-minute intervals at the GEOMAR pier in close distance from our setup (54°19'48.8" N 10°08'59.6" E) (AANDERAA oxygen sensor 3835 & SEABIRD SBE 37-SI MicroCAT CT(D)). The sensor system is mounted to a floating platform so that a continuous depth of 1m is guaranteed at every time point.

**Microalgae enumeration**

One-eight of each PB was cut and inspected under epifluorescence microscope (ZEISS Axio Scope.A1) for chloroplast enumeration at each time point. Considering the visibly patchy distribution of microalgae community on filters, we tried to cut a slice that could be a fair representative of distribution. Five different areas (replicates) from each slice were enumerated using imageJ software and the average number was reported for each PB or control sample. The area counted under the microscope was calculated as 2.327 square $mm^2$.

**16S rRNA gene sequencing and bioinformatic analysis details**

The two-step PCR strategy from Gohl et al. [2], the KAPA HIFI HotStart polymerase (Roche) and the Nextera XT v2 indexing primers were used to prepare amplicon libraries. Gel pictures were utilized for estimation and normalization of amplicon concentrations from the second PCR and amplicons were pooled accordingly. The primer dimer was removed through gel extraction (D4001; ZYMO-Research, Irvine, CA, USA). The final purified library was analysed by qPCR on a StepOnePlus™ Real- Time PCR System (Applied Biosystems), and bioanalyzer to assess the final concentration, and sequenced at the Max-Plank-Institute for Evolutionary Biology (Plön,

Germany). The sequenced library contained three blank technical controls from the DNA extractions (sterile beads, sterile filter, and sterile seawater were controlled for contamination), one negative PCR control and one positive control.

All bioinformatic processing were performed on the high-performance computing cluster (Hybrid NEC HPC system, University of Kiel, CAU) using QIIME 2 2022.11 [3]. FASTQ files were demultiplexed (Illumina bcl2fastq v2.20.0.422). After classification of ASVs against the SILVA 138.1 database. All ASVs were aligned with mafft [4] (via q2-alignment) and used to construct a phylogeny with fasttree2 [5] (via q2-phylogeny). Decontamination analysis was performed using "prevalence" method and the function "isContaminant" from the R package Decontam v1.18.0 [6] which selects contaminant ASVs based on the prevalence (presence/absence across samples) of each sequence feature in true positive samples compared to the prevalence in negative controls. This method was followed by a manual inspection for those ASVs present in seawater samples with lower number of replicates.

**Curation of Unassigned ASVs**

We used a phylogenetic approach for reclassification of Unassigned ASVs that can help with reclassification of mitochondria and chloroplast and detection of novel microorganisms not identified through typical search against databases. Our classification workflow assigned 313 out of 1409 sequences as Unassigned making up a total of 22% reads. Because these sequences could potentially be divergent prokaryotic sequences, we used a phylogenetic approach to evaluate and reclassify where possible. Initial observations via BLAST searches of NCBI revealed most

Unassigned sequences were mitochondria, therefore we focused a phylogenetics-based analysis on mitochondria and Rickettsiales, along with a less intense focus on a subset of diverse bacteria, and chloroplast sequences. To generate a reference alignment and tree, we selected all Rickettsiales sequences (which includes mitochondria), five sequences from all orders of Alphaproteobacteria and the 60 most common phyla in SILVA 138 database.  Sequences shorter than 1200 bp were removed and the remaining were clustered using cdhit (cd-hit-est v.4.8.1)[7] at 99% sequence similarity. Reference sequences were aligned using mafft software (v.7.505) with default settings. ASVs classified as Unassigned, Rickettsia, Mitochondria and Chloroplasts were then added to the alignment with mafft (via settings --add and –keeplength). Positions with greater than 50% gaps were removed with TrimAL (v.1.4.1). A phylogenetic tree was constructed using IQ-TREE (v.2.2.3) [8] by employing the SYM+ASC+R10 model parameters (selected based on substitution model selection within IQ-TREE [9]). Branch supports were obtained with the ultrafast bootstrap (UFBoot) [10] via 1,000 replications. The tree was visualized with FigTree (version v1.4.4) [11]. The tree was rooted at a long branch of Tistrellaceae (KC119130.1.1436), since no quey sequences were near to the branch. Unassigned ASVs which were clustered with mitochondria (n=295) and chloroplast (n=1) were considered as such and thus omitted from further analyses, while those which were confidently placed among prokaryotic phyla based on bootstrap support (>0.9) and branch length were classified as bacteria (n=14, making up 0.1 % of total reads) and were considered in downstream analyses. Unassigned ASVs with extremely long branch lengths were still considered unassigned (n=2) (Tab. S1-A, S1-B).

**Network construction and data preprocessing**

SPIEC-EASI was used for network inference based on meinshausen-buhlmann (MB) neighbourhood selection [12] on merged ASV datasets of prokaryotes and microalgae . Combining different types of datasets may require distinct processing steps depending on the characteristics of each microbial community, for instance variations in different domains abundances. This enables preserving the integrity of values within each domain while making them compatible. In the current study, considering higher numbers of microalgae reads on non-living substrate compared to seaweeds' biofilm, datasets of microalgae, after deletion of samples with below 100 reads (3 samples), were first rarefied at their minimum read depth for seaweeds (237) and proxy biofilms and controls (1114) separately, and then merged to the corresponding set of prokaryotic datasets (rarefied at their minimum read depth 6121 accordingly) in order to make a comprehensive microbial dataset for each group (in the format of phyloseq object). Only samples who had both eukaryotic and prokaryotic counterparts after rarefaction were considered for merged dataset (30 seaweed samples and 47 PBs and control samples).

We employed SPIEC-EASI as the technique for microbial association and correlation inference, where nodes and edges represent microbes and statistically significant associations between them correspondingly, to explore microbial associations and abundance correlations. This method was chosen since it considers the compositionality and sparsity of microbiome data (Kurtz et al., 2015). Clr transformation of data, sparse inverse covariance estimation and model selection are intrinsically implemented in spiec.easi function. For more finely sampling of the lambda path and obtaining a denser network, nlambda for each model selection was set to obtain at least 0.49 (closest possible value to the target stability threshold of 0.05). Ultimately, using SPIEC-EASI we could detect associations based on a (linear) relationship between ASV abundances whenever two ASVs are not conditionally independent. Data processing and networks construction were performed using R (version 4.1.1) combined with multiple R packages including phyloseq, SpiecEasi, igraph and microbiomeutilities and their dependencies. The module/submodule detection and modularity analyses were performed using fast greedy modularity optimization as a function in igraph [13]. Also graph variables such as natural connectivity (derived mathematically from the graph spectrum as an average eigenvalue), modularity, betweenness centrality, edge density (the ratio of the number of edges to the number of possible edges), edge betweenness, mean degree (i.e., the count of edges a node has) as a measure of sparsity, mean harmonic centrality, edge density, transitivity, and degree heterogeneity were obtained via igraph package in R. Network visualizations were done by importing the graphs data in Cytoscape (v. 3.9.1) and using Edge-weighted Spring Embedded Layout. Other data associated with ASVs, such as taxonomy and ASV tables, relative abundance, edge information, and positive and negative correlations, among- and between-module connectivity values were additionally imported in Cytoscape and visualized in network figures. Visualizations were enhanced using Adobe Illustrator (2023).

**SUPPLEMENTARY FIGURES**

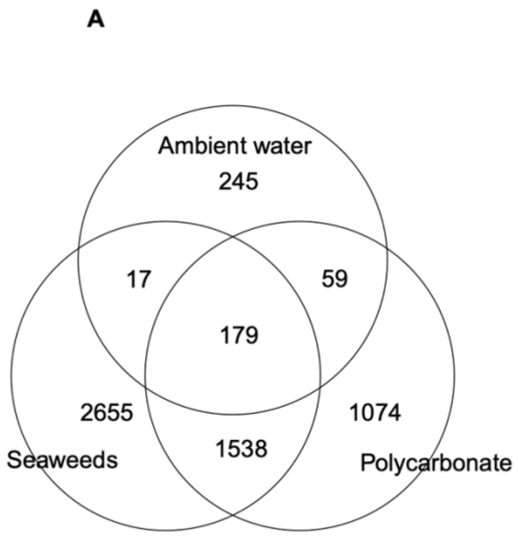


Fig.S1. Number of unique and shared prokaryotic (A) and microalgae ASVs (B) recovered from different substrates and ambient water.

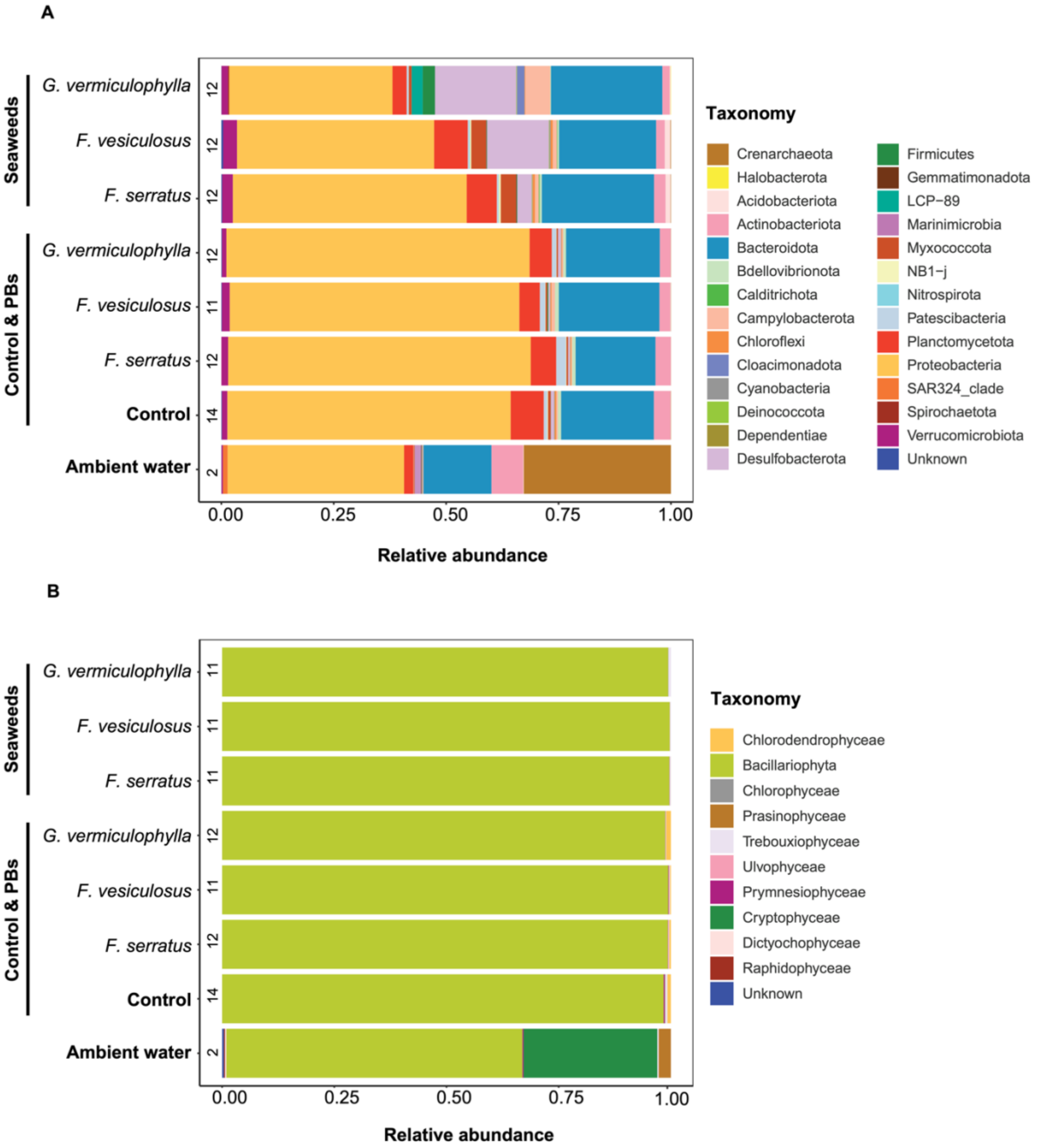


Fig.S2. Taxonomic composition of the microbial community, A) prokaryotes and B) microalgae, associated with seawater, proxy biofilms and control samples during two timepoints. Size of the stacked bars are proportional to relative abundances averaged within the replicates of the same group. The number of contributing samples for each group stated on the lefthand side of each bar.

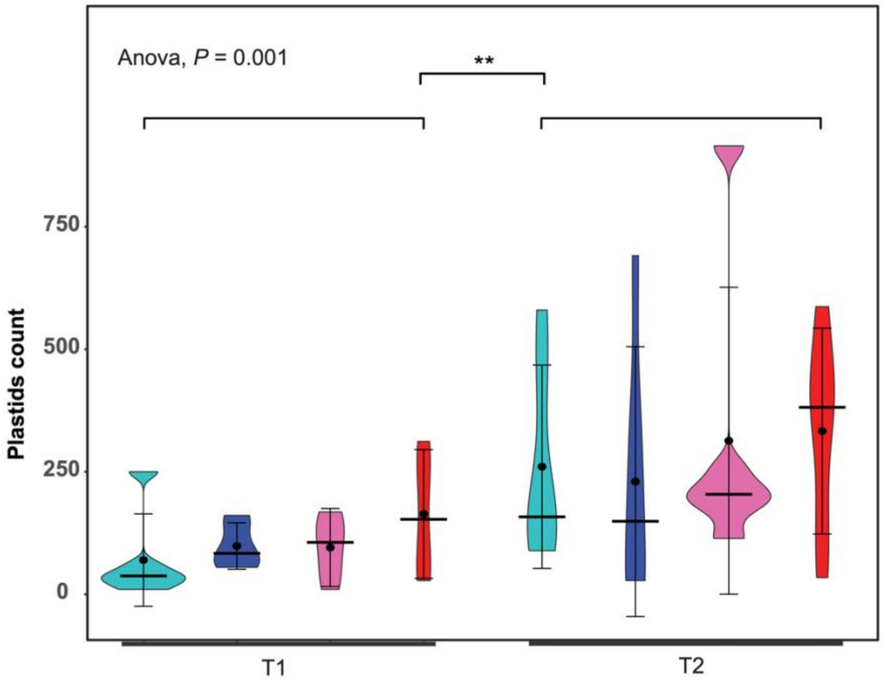


Fig.S3. Violin plots showing the average number of plastids count per area (2.327 $mm^2$) of proxy biofilms and control filters over two experiments. The plots show the median, interquartile range (including outliers) and the point and the bar showing the mean and the standard deviation. Six replicates were analyzed per timepoint for each treatment, with five areas counted per replicate.

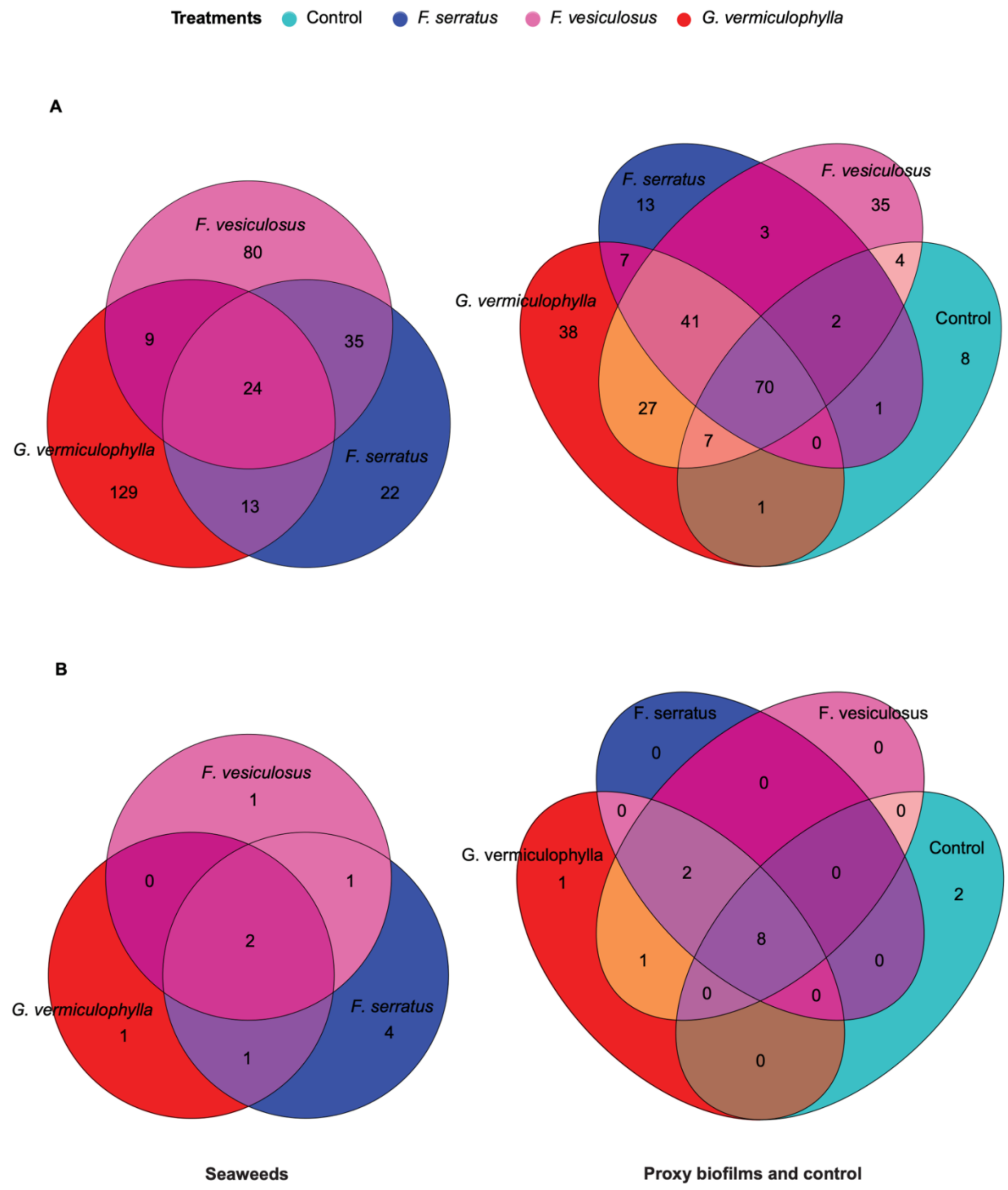


Fig.S4. Venn diagrams depicting unique and shared A) prokaryotic and B) microalgae ASVs among core microbiomes of seaweeds, proxy biofilms and control filters.

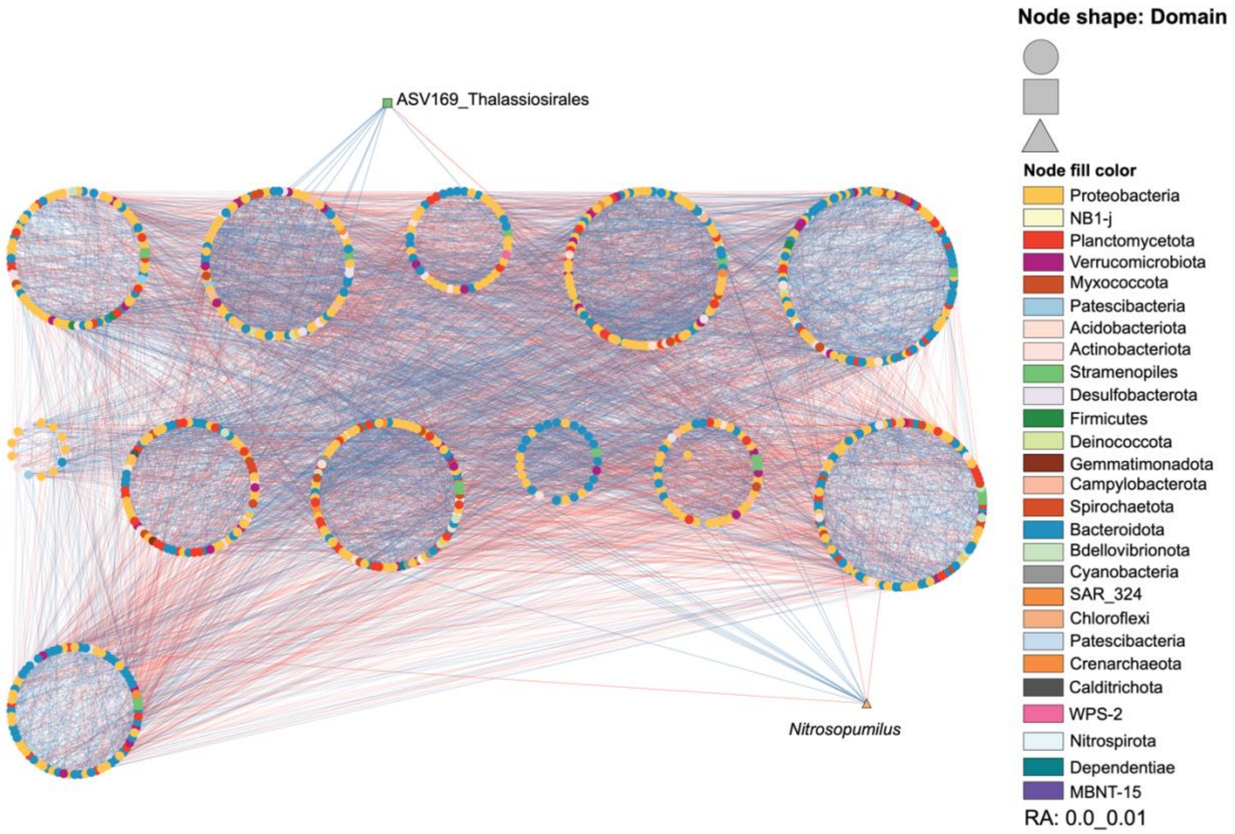


Fig.S5-A. WMC network of *Fucus serratus* (modular layout). Nodes represent ASVs and are shaped according to the taxonomic domains. Edge colour denotes a positive (blue) or negative (red) association between two connected ASVs with the width proportional to weight or strength of connection.

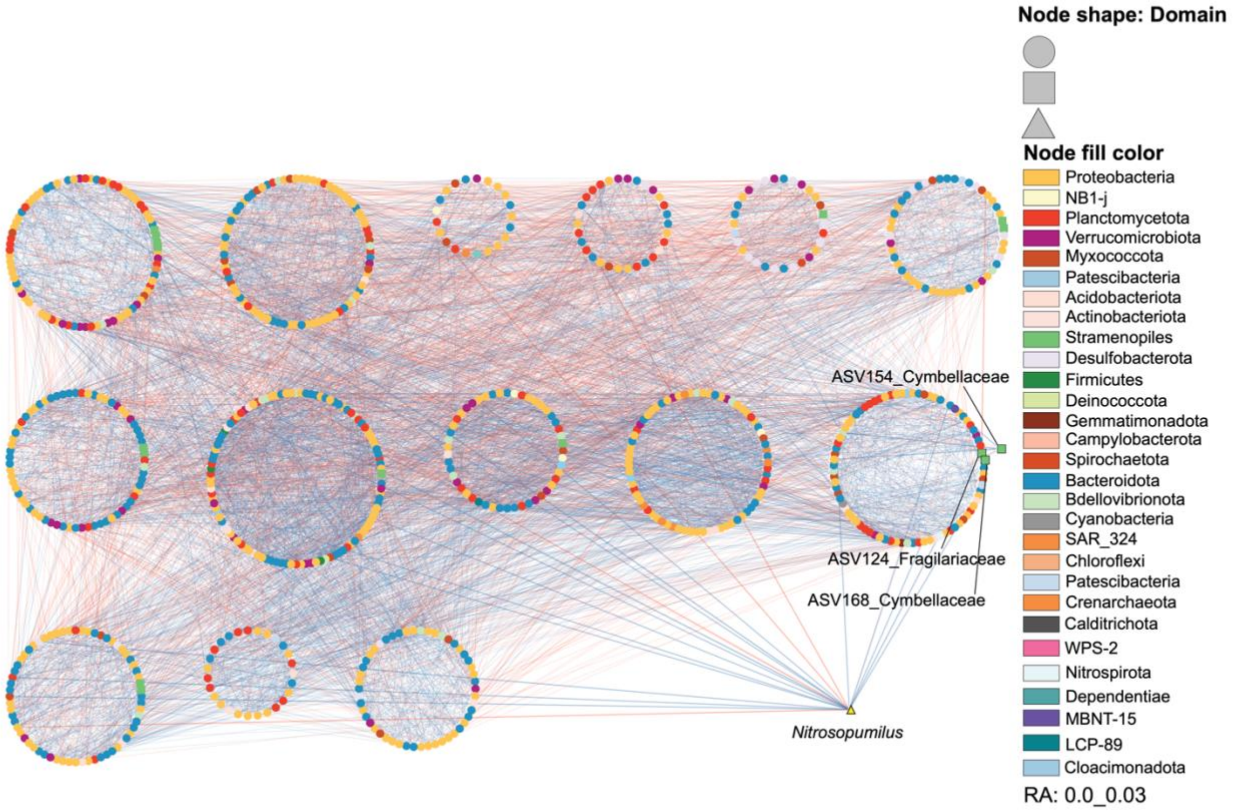


Fig.S5-B. WMC network of *Fucus vesiculosus* (modular layout).

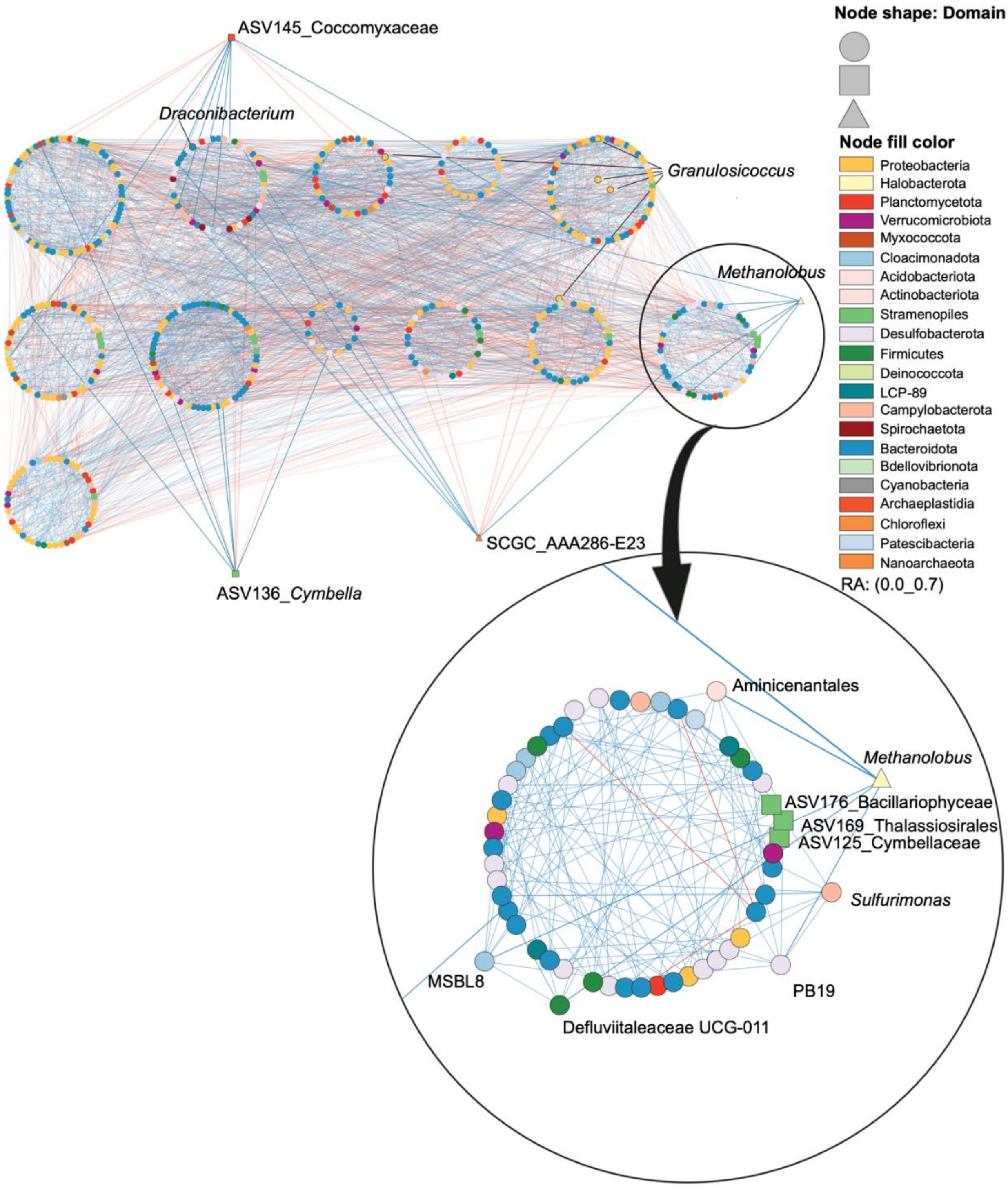


Fig.S5-C. WMC network of *Gracilaria vermiculophylla* (modular layout). Enlarged module shows the positive association (blue edges) between sulfur cycling bacteria (Desulfobacterota),

Aminicenantales (Acidobacteriota) [14], *Sulfurimonas* (Campylobacterota) [15], MSBL8 (*Cloacimonadota* ) [16], *Draconibacterium* (Bacteroidota) [17], and methanogenic archaea.

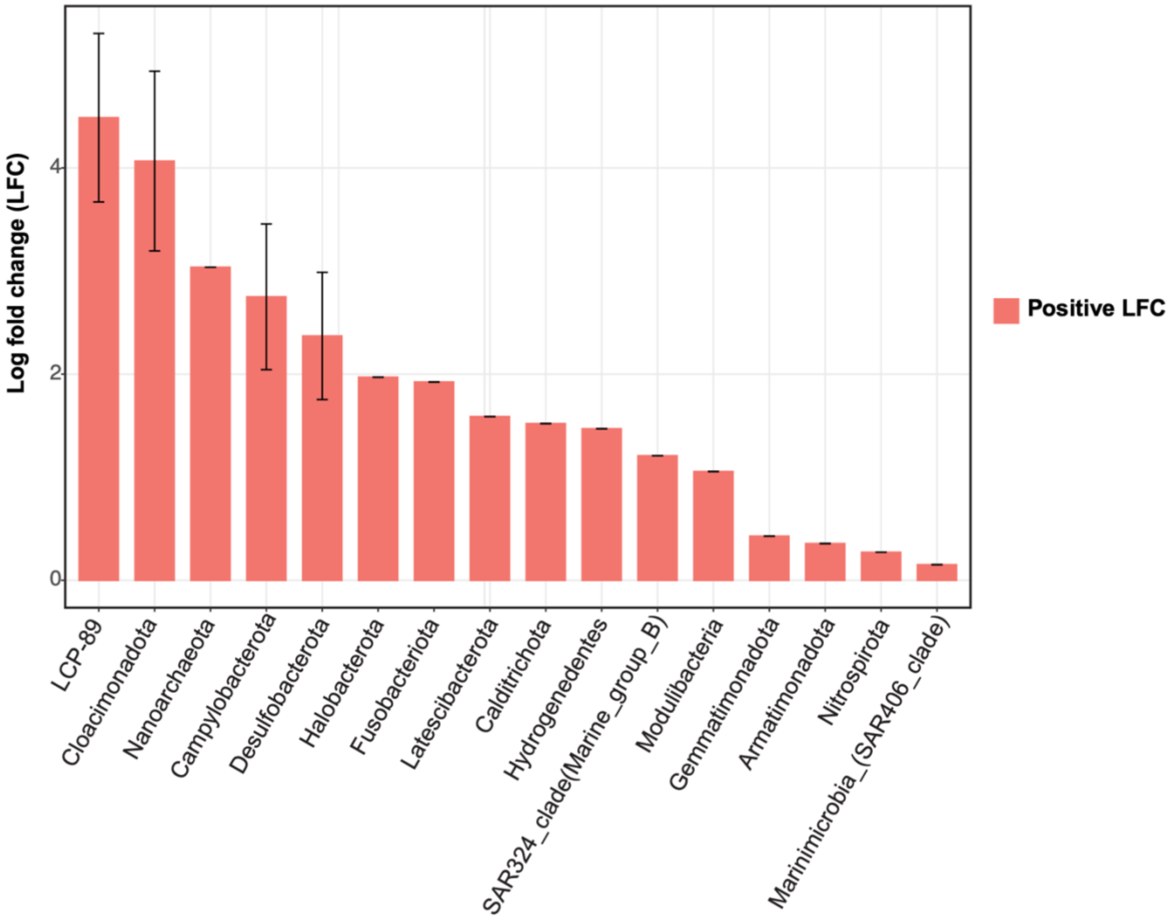


Fig.S6. Bar plot showing result from the ANCOM-BC log-linear model, determining taxa that were differentially abundant in the second timepoint compared to the first timepoint.

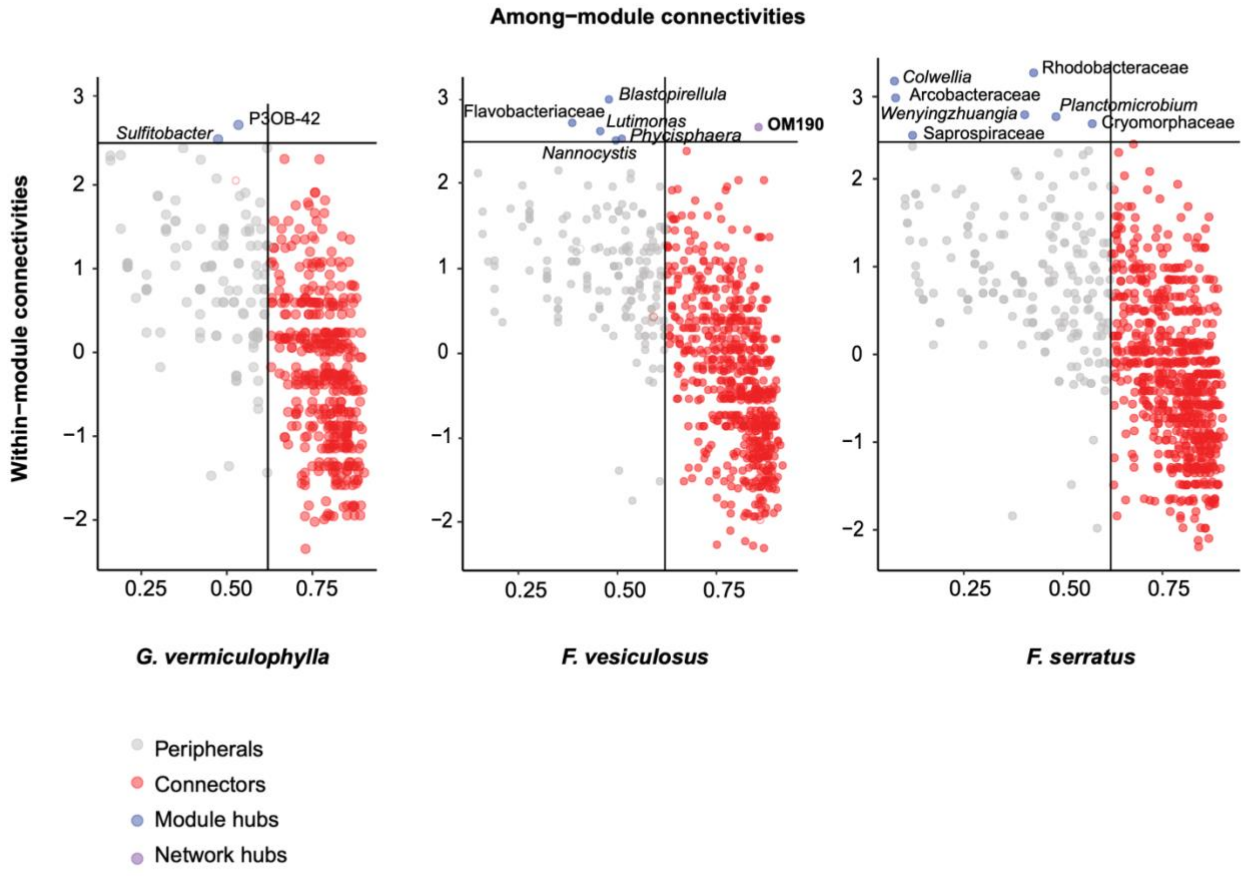


Fig.S7. Scatter plots showing distribution of microbial ASVs (prokaryotes and microalgae) according to their within-module and among-module connectivity. Each dot represents an ASV in the WMC dataset of each seaweed host. The four panels show the role distribution of selected groups of microbes. ASVs representing module, and network hubs are indicated on the plot.

### SUPPLEMENTARY TABLES

Tab.S1. Metadata file including algal weight, Shannon indices and environmental measurements.

Tab.S2. A) Reclassified ASVs after application of the phylogenetic approach for further classification of unassigned ASVs. B) Annotation file of phylogenetic tree used for reclassification of Unassigned ASVs.

Tab.S3. Statistical output of models fitted in this study. Diversity analysis (α-diversity, Shannon index) of A) prokaryotes on seaweeds and their paired PBs biofilm. B) prokaryotes on PBs and control filters. C) prokaryotes on seaweeds biofilm. D) microalgae on seaweeds and their paired PBs biofilm. E) microalgae on PBs and control filters. F) microalgae on seaweeds biofilm.

Tab.S4. Statistical output of models fitted in this study.   Enumeration of microalgae observed on PBs and control filters (average count provided based on 5 microscopic observation per filter_Tab.S12).

Tab.S5. PERMANOVA to compare A) prokaryotic community on a pair set of seaweeds and their PBs after removal of control (68 samples). B) prokaryotic community on all PBs and control filters. C) prokaryotic community on seaweeds biofilm composition. D) microalgae community on a pair set of seaweeds and their PBs after removal of control (68 samples). E) microalgae community on all PBs and control filters. F) microalgae community on seaweeds biofilm composition.

Tab.S6. RDA analysis assessing the impact of environmental factors on prokaryotic composition shifts in A) mature biofilm of *G. vermiculophylla*. B) *G. vermiculophylla* PBs C) all non-living substrates biofilms (PBS and controls).

Tab.S7-A) Diversity of prokaryotic core microbiome among different sample groups.
(SW: Seaweed Living substrate, PC: Polycarbonate non-living substrate. Two host-specific ASVs associated with *G. vermiculophylla* core microbiomes in both mature biofilm and PB are shown in yellow (corresponding to their Proteobacteria phylum colour code).

Tab.S7-B) Diversity of eukaryotic core microbiome among different sample groups. (SW: Seaweed Living substrate, PC: Polycarbonate non-living substrate).

Tab.S8. Network mean variables used for comparison of microbial connectivity between sample groups visualized via heatmap and PCA plots.

Tab.S9. Microbial association network variables for all nodes and their interactions among all sample groups.

Tab.S10. ASVs shared between mature biofilm and PBs for each seaweed species

Tab.S11. Hub taxa identified for different sample groups on living vs non-living substrates.

Tab.S12. Number of enumerated microalgae on PBs and control filters over two experiments. Count 1 to 5 represent 5 replicates per filter. Each replicate covered 2.327 square millimeter area of the filter.
